# Admixed Populations Improve Power for Variant Discovery and Portability in Genome-wide Association Studies

**DOI:** 10.1101/2021.03.09.434643

**Authors:** Meng Lin, Danny S. Park, Noah A. Zaitlen, Brenna M. Henn, Christopher R. Gignoux

## Abstract

Genome-wide association studies (GWAS) are primarily conducted in single-ancestry settings. The low transferability of results has limited our understanding of human genetic architecture across a range of complex traits. In contrast to homogeneous populations, admixed populations provide an opportunity to capture genetic architecture contributed from multiple source populations and thus improve statistical power. Here, we provide a mechanistic simulation framework to investigate the statistical power and transferability of GWAS under directional polygenic selection or varying divergence. We focus on a two-way admixed population and show that GWAS in admixed populations can be enriched for power in discovery by up to 2-fold compared to the ancestral populations under similar sample size. Moreover, higher accuracy of cross-population polygenic score estimates is also observed if variants and weights are trained in the admixed group rather than in the ancestral groups. Common variant associations are also more likely to replicate if first discovered in the admixed group and then transferred to an ancestral population, than the other way around (across 50 iterations with 1,000 causal SNPs, training on 10,000 individuals, testing on 1,000 in each population, p=3.78e-6, 6.19e-101, ~0 for F_ST_ = 0.2, 0.5, 0.8, respectively). While some of these F_ST_ values may appear extreme, we demonstrate that they are found across the entire phenome in the GWAS catalog. This framework demonstrates that investigation of admixed populations harbors significant advantages over GWAS in single-ancestry cohorts for uncovering the genetic architecture of traits and will improve downstream applications such as personalized medicine across diverse populations.

## Introduction

Genome-wide association studies (GWAS) have allowed for significant progress in the field of human complex traits. However, groups with multiple ancestral origins have seldom been a primary focus in large scale genetic studies because: (1) admixed groups, along with other non-European populations, have largely been underrepresented in GWAS designs in the past (Bustamante et al., 2011; Popejoy and Fullerton, 2016; Martin et al., 2017a), and (2) the population structure from heterogeneous ancestries in an admixed group, if not properly corrected, can result in spurious correlation signals and thus greater false positive rates (Rosenberg et al., 2010). However this mixture of ancestries present in admixed populations provides opportunities for novel discovery. Recent advancements in methodologies tailored for genetic mapping in admixed populations include disentangling of ancestry principal components and relatedness in the presence of admixture (Thornton et al., 2012; Conomos et al., 2015, 2016), combining local ancestry and allelic information to improve quantitative trait locus (QTL) mapping (Pasaniuc et al., 2011; Shriner et al., 2011; Atkinson et al., 2021), leveraging local ancestries for detection of epistasis (Aschard et al., 2015), and better fine mapping from linkage disequilibrium (LD) variability in diverse groups (Zaitlen et al., 2010; Asimit et al., 2016; Wojcik et al., 2019; Shi et al., 2020). Despite the fast development and practicality of these methods, they have not often been applied to sample sizes of hundreds of thousands to millions because study design and data collection in mega-scale cohorts routinely prioritize recruitment of participants of single ancestry (Atkinson et al., 2021).This greatly impedes downstream progress, such as polygenic risk score application across populations, where much lower accuracy is observed in non-European populations for many traits (Duncan et al., 2019; Martin et al., 2019; Cavazos and Witte, 2021).

In addition, complex traits in admixed groups potentially harbor differing genetic architectures and varying environmental exposures compared to most widely studied groups such as Europeans. Some biomedical traits have higher risk prevalence in admixed groups, such as prostate cancer in African Americans (Bhardwaj et al., 2017; Conti et al., 2021), asthma in Puerto Ricans (Lara et al., 2006; Pino-Yanes et al., 2015), obesity and type II diabetes in Native Hawaiians (Maskarinec et al., 2009), and active tuberculosis in a South African admixed population (Chimusa et al., 2014), which are likely attributed to elevated ancestry-specific risk allele frequency. Among anthropometric traits, skin pigmentation in groups with admixed ancestry harbor greater phenotypic variance than those with single ancestries (Martin et al., 2017b). Here, the larger phenotypic variance is likely caused by increased polygenicity in admixed groups, where in contrast some causal variants are nearly fixed in the single ancestry groups due strong directional selection of skin pigmentation (e.g., rs1426654 in *SLC24A5* (Lin et al., 2018)). The increase in minor allele frequencies in admixed populations compared to the populations of ancestral origin could be ubiquitous in traits that have been under differential processes of selection among ancestral populations or simply among populations that are deeply diverged. This would theoretically result in greater power of discovery in GWAS, as the analysis is most powerful for variants with higher minor allele frequency.

While genetic epidemiologists have typically focused on homogeneous populations, there are clear opportunities to improve discovery in admixed populations. For example, local ancestry can be leveraged to improve power in certain scenarios (e.g. (Pasaniuc et al., 2011)). In addition, Zhang and Stram observed a power gain in admixed individuals in dichotomous traits compared to pooled ancestral populations with stratification without environmental confounding (Zhang and Stram, 2014). Here, we extend a similar framework using a flexible mechanistic model to address the question of power compared to a similar-sized each ancestral population on its own, across a range of allelic differentiation (as measured by F_ST_ (Weir and Cockerham, 1984))with varying narrow-sense heritability, and across a range of ancestry–phenotype associations, whether driven by genetics or environment. We further extend these insights to look at power for replication, whether from admixed populations to ancestral populations or *vice versa*, as well as opportunities to derive trans-ethnic polygenic scores.

## Methods

### Simulation-based power estimate between a trait and global ancestry

The first simulator we provide in this study builds phenotypes in admixed populations using only global ancestries, without involving genotype. The aim is to assess if the sample size is adequate for observing a dichotomous trait by ancestry correlation. The details are described in *Supplementary Notes*.

### Genotype-mediated simulation framework

The general simulation framework consists of two steps: first, we model ancestries and simulate genotypes based on ancestry specific frequencies and phenotypes (Fig. 1); then, we test associations between the phenotype and causal variants via a linear model for a quantitative trait, or a logistic regression for a dichotomous trait, and summarize the statistical power. If the population is admixed, global ancestry is supplied as a fixed effect to correct for population structure.

**Figure 1.**
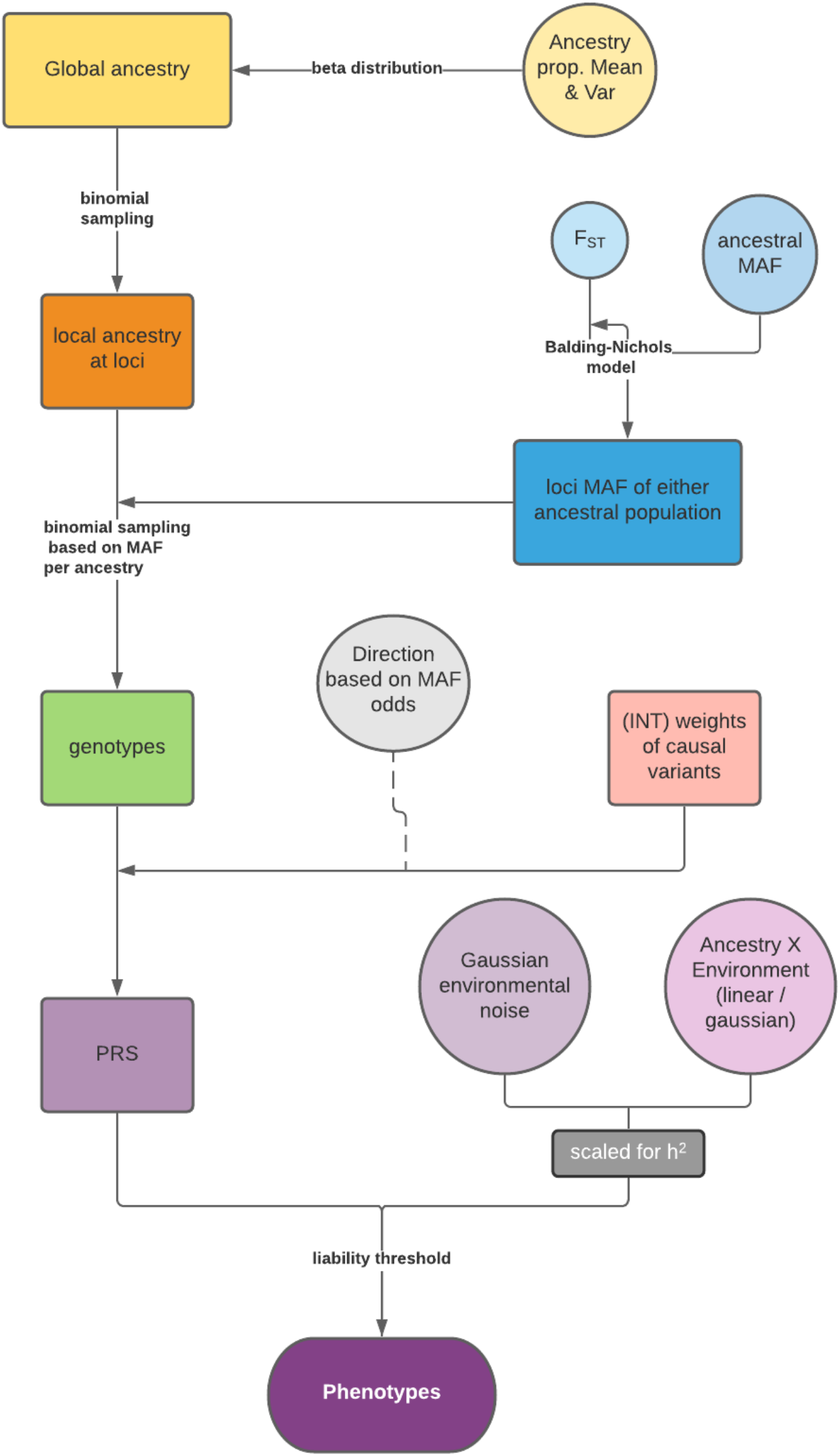
Flow chart of the genotype-mediated simulation framework (prior to association testing).

#### 1) Ancestry modeling

Global ancestry in a 2-way admixed population is modeled as a beta distribution. The *i*th individual’s ancestry *θ* is characterized as

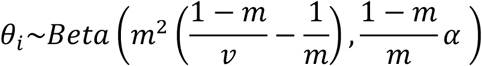

where *m* and *v* are the mean and variance of the global ancestry (from a presumptive population 1 in this framework) in an admixed population of interest.

The local ancestries, i.e. the source of ancestry of both the maternal and paternal copies of haplotypes at any genomic position, can be obtained from a binomial sampling with the probability equaling the global ancestry. The process is repeated independently for diploid chromosomes over a presumptive number (*n*) of LD-independent loci to form a *n* × *N* local ancestry matrix, where N is the proposed sample size.

#### 2) Genotype simulation

We first draw allele frequencies in the two ancestries (Population 1 and 2) from a beta distribution under the Balding–Nichols model (Balding and Nichols, 1995) with a given F_ST_

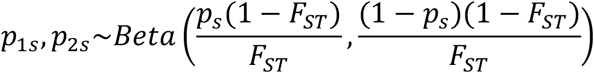

where *p_s_* is the allele frequency at an independent locus *s* in an ancestral population prior to the divergence and drawn from a uniform distribution *unif*(0.001, 0.999). Within the model we additionally provide additional distributions if the focus is not on common variants as it is here. We set F_ST_ as a flexible value to increase from a baseline genome-wide F_ST_ when the ancestral allele becomes rare. This is to reflect that a rarer variant in the ancestral population is easier to drift to different frequencies in diverged populations, especially if one population has undergone a severe bottleneck (as would be expected to increase F_ST_).

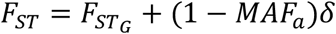

where 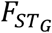 is the genome-wide background F_ST_ between the two populations of ancestral origin and is considered lower than the F_ST_ between trait-causal loci because of the difference in directional selection. *MAF_a_* is the minor allele frequency of the variant in ancestral populations and δ is the increment with regard to minor allele frequency (MAF) decrease, set as 0.3 in this study. Alternatively, we also test for fixed *F_ST_* under the genome-wide background value when exploring the effect on power from various F_ST_ values ranging from 0.1 to 0.9. The genotypes are then drawn from binomial sampling using the allele frequency corresponding to the local ancestry assigned at the locus (i.e., *p*_1*s*_ or *p*_2*s*_) across all loci.

#### 3) Genetic contribution to trait

We randomly assign *w* out of the total *n* loci to be causal variants, where *w* is the proposed polygenicity of the trait. The weights for the causal variants are drawn from a standard normal distribution *N*(0,1), and the signs of the weights are tied to the prevalence of the allele in the two populations of ancestral origin: the direction of the weight, positive or negative, at a locus is decided by the binary outcome of trial with probability 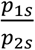. In this way, a difference in directional selection of the complex trait in the two populations is introduced to facilitate a correlation between the trait and ancestry in the admixed group. Then, polygenic risk scores (PRS) in samples can be calculated based on the weights and the genotypes at causal loci.

#### 4) Non-genetic contribution to trait

The non-genetic component is treated as the sum of two parts in admixed populations: (1) random environmental variation modeled as Gaussian noise and (2) environmental confounders correlated with ancestry, such as socioeconomic status and education, modeled as ancestry by environment interaction. Details are described in *Supplementary Notes*.

#### 5) Phenotype

For quantitative traits, the phenotype is the direct sum of the genetic component (i.e. PRS) and the non-genetic score. For dichotomous traits, the phenotype is converted from the sum of genetic and non-genetic scores to binary case and control status based on the given liability threshold of the case prevalence.

#### 6) Association testing

Association between the trait and a variant is tested via a linear regression for a quantitative trait, or a logistic regression for a dichotomous trait in all three populations. The global ancestry is corrected in the admixed group. Power is defined as the proportions of causal variants with a significant p value above a given stringency threshold.

### Estimation of false positives

A false positive rate in associations is empirically verified against the association stringency by calculating the proportions of non-causal variants being discovered with significant association p values. This is calculated separately in each population.

### PRS estimation

For training purposes, we obtained the “estimated” weights of causal variants by conducting association analyses in 10,000 individuals in each population. We then used these weights to estimate PRS in another 1,000 individuals in each population as a test. We tested the PRS construction in two ways: firstly, we only used significant (p<0.05) causal-variants from the training set (true positives); secondly, we included all significant variants (all positives) over a range of different stringency (p=0.05, 5e-4, 5e-6, 5e-8, respectively). The true PRS of the individuals in the test set were available through an intermediate step in the simulations (Fig. 1) and were used to test the accuracy of the estimated PRS via correlation coefficients.

### Calculation of F_ST_ for traits from the GWAS catalog

We used the full NHGRI-EBI GWAS catalog “All associations v1.0” (Buniello et al., 2018) to extract variants that are significant genome-wide (< 5e-8). We restricted traits to 899 that have more than 10 significantly associated variants that can be found in the 1000 Genome Project Phase 3 (Consortium et al., 2015), and computed Weir and Cockerham’s F_ST_ (Weir and Cockerham, 1984) between 99 Utah Residents (CEPH) with Northern and Western European Ancestry (CEU) and 108 Yorubans in Ibadan, Nigeria (YRI) samples using PLINK v1.9 (http://www.cog-genomics.org/plink/1.9/) (Chang et al., 2015). The genomic background weighted F_ST_ was calculated on common variants (MAF >5%) only.

## Results

### Correlation between a trait and ancestry is common

Complex trait studies in groups with heterogeneous ancestries usually require a correction for population structure. The implicit assumption is often a correlation between global ancestry and the trait that is commonly observed *a priori*. The estimated ancestries, or typically ancestry informative principal components, are included as a fixed effect to adjust for phenotypic variance from non-genetic confounders (e.g., social and cultural factors correlated with population structure), and to avoid spurious associations (Price et al., 2006). The correlations between ancestries and complex traits can also be due to changes in genetic architectures among ancestral groups either due to differential selection or deep divergence among populations. This in turn forms one of the basic motivations of multi-ancestry genetic studies, including admixture mapping (loci with ancestry deviating from genome-wide expectation), and cross-population transferability of genetic predictors. Therefore, we provide a power estimate for whether a significant correlation with ancestries can be observed, within a given incidence rate and ancestry distributions (Methods, Fig. S1).

### The power of genetic discovery in an admixed population is higher than in ancestral populations

We primarily focused on a genotype-mediated simulation framework to investigate the GWAS setting in an admixed group. We started by modeling global ancestries, then generating LD-independent genotypes based on population divergence, and subsequently the corresponding phenotypes under an additive model (Methods, Fig. 1). We set up the model in a 2-way admixed group with similar proportions to African Americans, here an average of ~75% West African ancestry (denoted as Population 1 in simulations) and ~25% European ancestry (denoted as Population 2) (Bryc et al., 2015; Baharian et al., 2016). We simulated a complex trait assuming 50% narrow sense heritability with 100 causal variants, either as a quantitative or a dichotomous trait with a liability threshold of 5%, in both the admixed population of interest and the homogenous populations of ancestral origin (N=1,000 each). To induce a difference between ancestral phenotypic distributions and a correlation between a trait and global ancestries, we tied the direction of effect sizes to the minor allele frequencies in the two populations of ancestral origin (Methods, Fig. 2). In this study, each independent setting was repeated in 50 runs.

**Figure 2.**
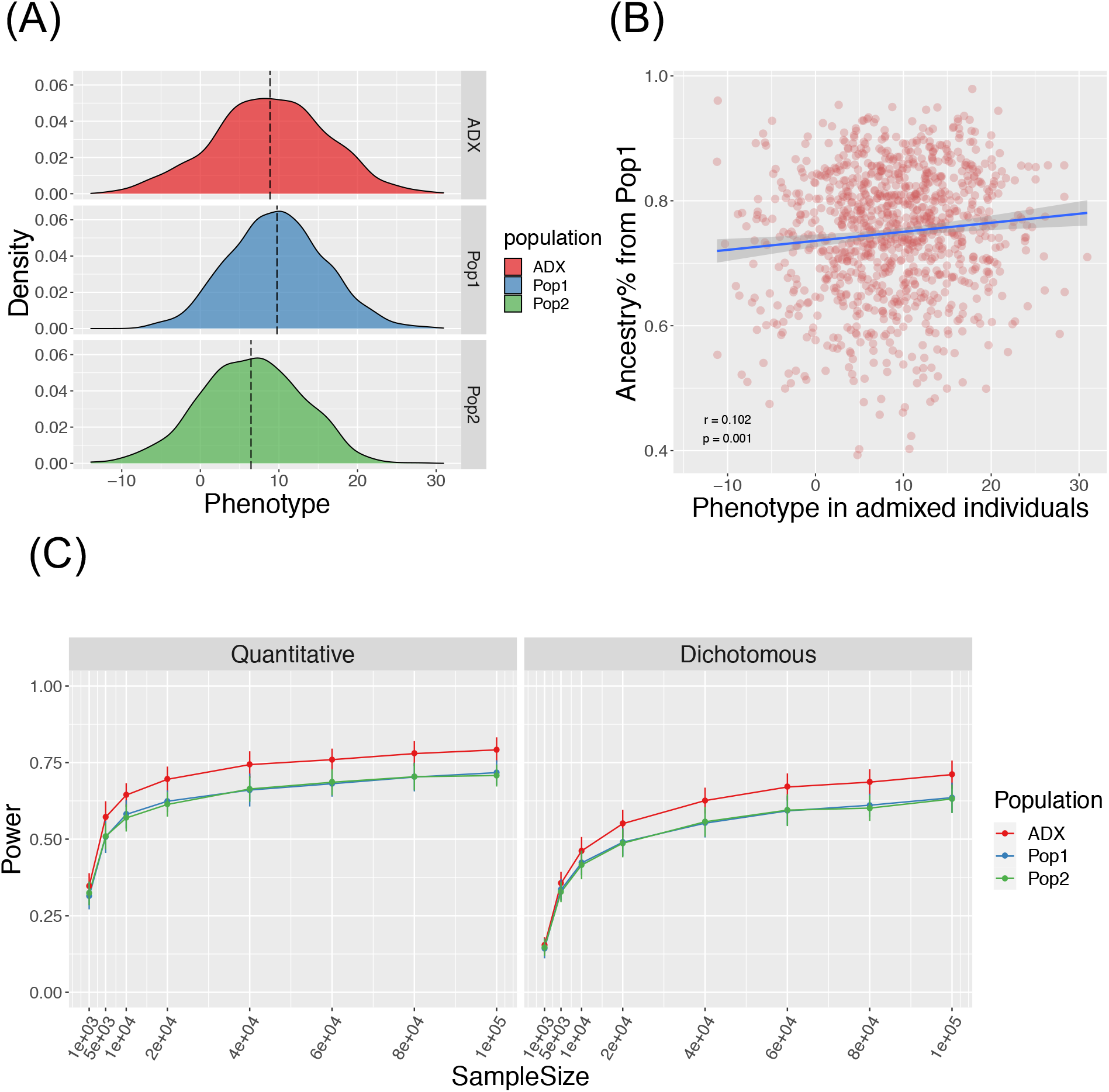
Genotype mediated simulation under an example condition. The simulated trait has 100 causal variants with a narrow sense h^2^=0.5, F_ST_ at causal variants 0.2+ (explained in Result), and environment by ancestry effect modeled as the sum of ancestral Gaussian environmental noise proportional to global ancestry. (A) Simulated quantitative phenotype distribution of populations of ancestral origin (Pop1, Pop2, blue and green respectively) and admixed population (ADX, red) of 1,000 samples each. (B) Correlation between simulated phenotype in admixed population and the global ancestry from Population 1. (C) Power to discover a causal variant over a range of sample sizes in a quantitative and dichotomous trait. Data point and error bars represent the mean and standard deviation across 50 repeats, respectively.

In the standard setting, we modeled the parameters described above, and F_ST_ across the 100 causal variants between Population 1 and 2 as a flexible value with a baseline equaling the genome-wide F_ST_ of 0.2 and an increment associated with the rarity of the ancestral MAF. This is referred to as Fst=0.2+ in the text. The aim of this is to mirror the larger stochasticity in the frequency change of an ancestrally-rare variant in diverged populations, especially when one of the derived populations has experienced the severe out-of-Africa bottleneck. We then tested associations between the phenotype and each locus while correcting for global ancestries over various sample sizes ranging from 1,000 to 1,000,000. We found the power to discover a causal variant at a canonical threshold of p ≤ 0.05 significantly higher in an admixed population than the average in either of the populations of ancestral origin (Wilcoxon p=1.23e-20 and 5.79e-10 across the range of sample sizes for quantitative and dichotomous traits described in Fig. 2, respectively).

In addition to the standard setting where an environment by ancestry effect (Env × Anc) is modeled as ancestry-weighted Gaussian noise, we explored an alternative where we model Env × Anc as linearly dependent on the ancestry percentages, which would explain a range of proportions of phenotypic variance from 0% to *1-h^2^* (Fig. S2). The power advantage in admixed populations remains consistent between the default Gaussian Env × Anc and linear modeling, where the latter was set as up to 10% of non-genetic components (Fig. S3).

The comparatively high power in admixed populations is more pronounced when the trait distributions have greater distance between Population 1 and 2, or the two populations are more deeply diverged, reflected by the larger F_ST_ at causal variants (Fig. 3). To relate to real-world GWAS, we compared our levels of differentiation to the NHGRI-EBI GWAS catalog (Buniello et al., 2018). Among the 899 traits that have more than 10 genome-wide significant hits found in 1000 Genomes Project, the majority (N=877) have at least one associated variant beyond the background F_ST_ of 0.155 (Fig. 3), we provide a list of the most-differentiated traits between CEU and YRI in Table S1. In contrast to the response to varying F_ST_, the statistical power does not obviously change when the narrow sense heritability of the trait differs (Fig. S4). When increasing the overall stringency of the type I error rate up to a conventional genome-wide significance of 5e-8, the power advantage remains very similar across different thresholds, despite the expected decrease in power value on the absolute scale in all populations (Fig. S5, S6). Thus we picked the canonical threshold of p ≤ 0.05 for the remaining analyses, as it can represent all stringency levels when this study focuses on the relative power comparison, and this more relaxed cutoff would include a larger number of causal variants for further discussion. The actual false positive rate of associations, as calculated from the 900 non-causal variants from the simulation, remained at approximately 5% in admixed samples across the full F_ST_ and h^2^ range, though it was lower in Population 1 and 2 as F_ST_ increases high enough to drive minor allele frequencies towards zero (Fig. S7).

**Figure 3.**
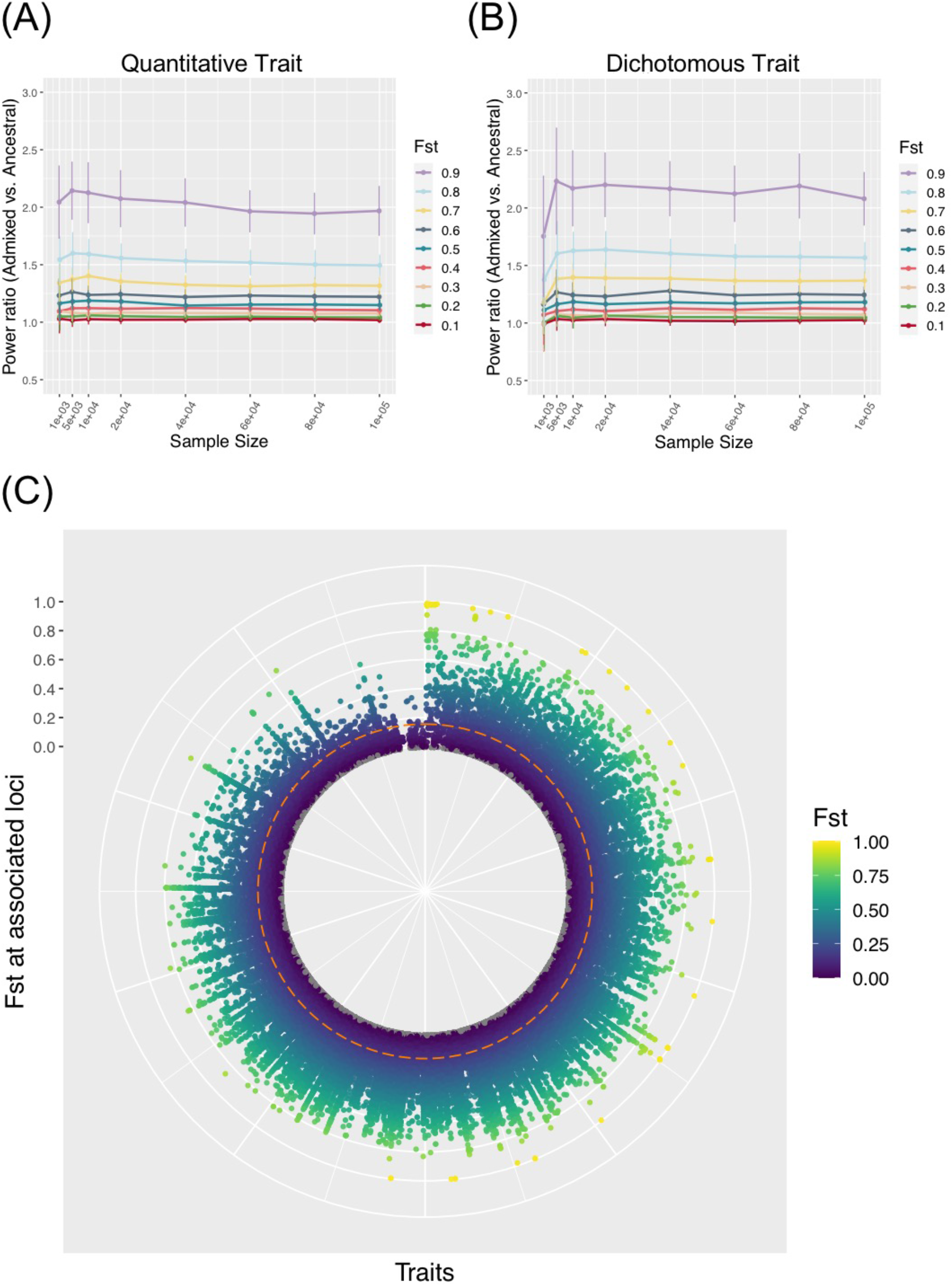
Varying F_ST_ at trait associated loci. Ratio of power in admixed population over the average in the two populations of ancestral origin, with different F_ST_ at causal loci in (A) a quantitative trait and (B) a dichotomous trait. F_ST_ was set to constant during simulations per a specified value. The trait was assumed to have 100 causal loci and a narrow sense heritability of 0.5, with environment by ancestry effect modeled as a sum of ancestral Gaussian noise proportional to the global ancestry. Data points and error bars represent the mean and standard deviation across 50 repeats, respectively. (C) F_ST_ at genome-wide significant hits for 899 traits from the GWAS catalog, between CEU and YRI from the 1000 Genomes Project Phase 3. Traits are spread along the radian (x-axis), with variant F_ST_ shown along the radius (y-axis). The dashed line represents the genomic background F_ST_.

### Cross-population replication and transferability is asymmetric between the admixed group and homogenous groups

As GWAS is conventionally focused on common variants, to investigate replication and transferability we then increased the polygenicity of a trait to 1,000 causal variants, and set MAF filtering at 5% for each population’s genotypes prior to testing associations. We compare discovery in the major ancestral population (Population 1) relative to the admixed population. The proportion of significant signals that replicate in the reciprocal group is asymmetric between the two populations. Discovery in the admixed samples was more likely to replicate in Population 1 than the other way around, and this trend becomes more exaggerated as trait F_ST_ increases (one-way Wilcoxon p=3.78e-6, <2.2e-16, and <2.2e-16 for F_ST_ = 0.2, 0.5, and 0.8, respectively; Fig. 4).

**Figure 4.**
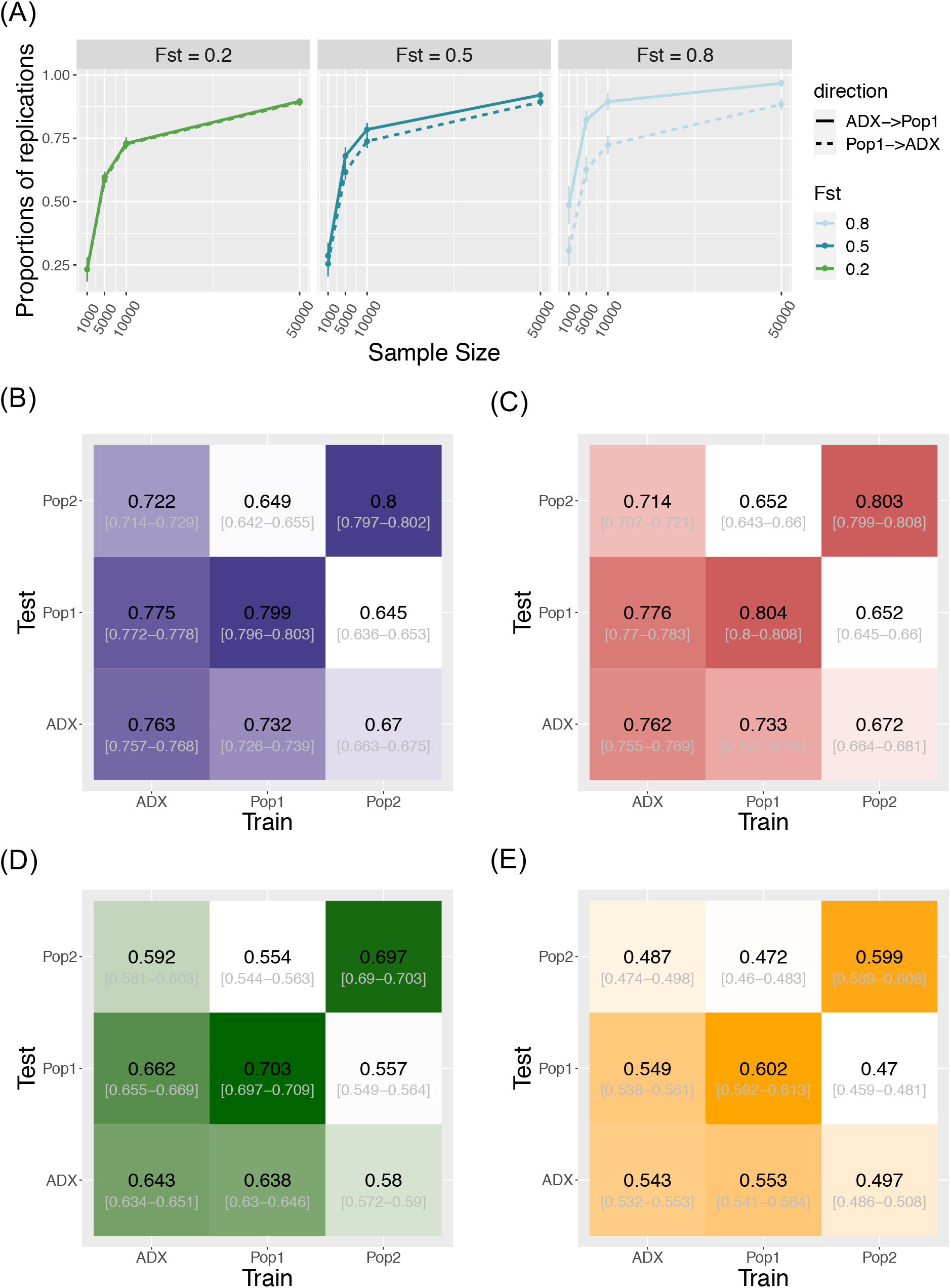
Transferability of GWAS variants across populations. (A) Replication of individual signals that are common in both ancestral Population 1 and the admixed group. Direction of replication is shown as a solid or dashed line: the former indicates loci are discovered in an admixed population and replicated in Population 1; the latter loci are discovered in Population 1 and replicated in the admixed population. Data point and error bars represent the mean and standard deviation across 50 repetitions. (B), (C), (D), and (E) Heat map of accuracy of PRS using signals above different stringency of significance level at 0.05, 5e-4, 5e-6, and 5e-8, respectively. The accuracy is measured as the correlation coefficient between the estimated PRS against the true PRS. The training population where the weights and variants were identified, and the test population in which to construct PRS, are specified on the x- and y-axes. Central numbers in black within each cell are the average correlation coefficient across 50 independent simulations, with the 95% confidence interval of the mean acquired from bootstrapping (n=1,000).

We tested the cross-population transferability of polygenic scores (or polygenic risk scores, PRS) constructed from discovered loci with MAF > 5% in each population by increasing the sample sizes of the training set to 10,000 per population and separately estimating the GWAS-based PRS in an additional testing set of 1,000 samples in each population. We measured the PRS accuracy as the correlation coefficient *r* between the estimated values and the true value in each test group across 50 repeats of simulations. Interestingly, the prediction accuracy is also asymmetric between admixed and homogenous samples. When we constructed PRS using only true positive signals at an alpha of 0.05, the accuracy of estimating PRS in Population 1 or 2 using weights and loci trained from the admixed population is significantly higher than the other way around. This holds true when using all (both true and false) positive signals at various stringency levels (Fig. 4; Figure S8; Table S2), suggesting another advantage in conducting GWAS in admixed populations.

## Discussion

Our simulation framework, APRICOT, Admixed Population poweR Inference Computed for phenOtypic Traits, demonstrated that GWAS in admixed populations has greater power for discovery than in the homogenous populations of ancestral origin, given the same sample sizes. The difference in power increases when the trait is under more differentiated polygenic selection in the two populations of ancestral origin, reflected by F_ST_. This is because when a trait is driven by more-differentiated variants, its causal variants are likely to be pushed to more extreme allele frequencies, thus weakening the statistical power of discovery in that population. In contrast, the frequency of the same causal loci in admixed populations likely have become more intermediate due to variation in ancestries, making them much easier to detect. An extreme yet classic example that echoes with the observation would be skin pigmentation, where selection is in the opposite direction in populations at high latitude and those living near the equator. A non-synonymous, skin-lightening mutation at rs1426654 is fixed in European descendants, with a high F_ST_ of 0.985 between CEU and YRI. This mutation would never have been discovered through GWAS if analyses were only conducted in European populations, but it is highly detectable through association analyses in admixed populations (Martin et al., 2017b).

Additionally, the power advantage in admixed populations may persist even for traits that have not been under such strong differentiation: for almost all the 899 traits we examined from the GWAS catalog, some associated SNPs can have a much larger than background F_ST_ between CEU and YRI, even when the traits themselves on average show limited differentiation (Fig. 3). We note however that the high F_ST_ across these trait-associated variants could partially be attributed to ascertainment bias, where the “tagging SNPs” by design are common in Europeans, making the corresponding genetic component of these traits seemingly more differentiated across populations (Novembre and Barton, 2018). The true causal variants that were tagged by these signals could have moderately attenuated F_ST_, yet the differences in allele frequency likely remain larger than expected, as previously observed from GWAS on simulated whole genome sequences between Africans and Europeans (Kim et al., 2018). Therefore, attempts to discover variants similar to these “F_ST_ outlier” signals would benefit from GWAS designed in admixed samples.

In this study, we provide a mechanistic framework to explore the relationship between power gain in single variant associations and variation in ancestries, mediated by the nature of intermediate allele frequencies in admixed populations. A similar hypothesis of power increase was also explored via simulations in Zhang and Stram (Zhang and Stram, 2014), though the focus of their study was to explore the role of local ancestry in genetic associations; therefore, the assumptions of architecture for comparison between admixed and ancestral populations were simplified, where non-genetic components (such as environmental effect and environment by ancestry interactions) and heritability were not considered in the model, and a constant effect size was assumed for all causal variants. Under this model, Zhang and Stram observed a power increase in admixed groups when compared to stratified analyses in the ancestral populations pooled with a proportion identical to the mean global ancestry percentage. Our simulations extended this framework to dive deeper into more-realistic scenarios across various ranges of environmental effect, trait divergence, and heritability. Moreover, we modeled the ancestry-phenotype association observed in many real-world traits under various different distributions. With a more adjustable genetic architecture in the model, we were able to quantify power advantage in admixed populations within different circumstances in order to investigate practical applications as replication portability and PRS.

In realistic practice, some confounders and restrictions beyond the model assumptions exist: first, some non-additive genetic components, such as genetic by environment interactions (G × E) and epistasis (Park et al., 2018; Rau et al., 2020), could potentially induce effect size heterogeneity at causal loci with or among populations (Rosenberg et al., 2019), thus obscuring the prediction of power advantage in admixed populations because the power of discovery would be variant-specific and balanced by the gain *vs*. loss from the increase in frequency and change in effect size. However, the increase in power is still expected to be substantial from additive components that are usually considered major in a genetic architecture, with effect sizes highly similar across populations (Wojcik et al., 2019). Additionally, currently the contribution from epistasis or G × E components to most trait variability is estimated to be relatively small (Wang et al., 2019; Dahl et al., 2020; Hivert et al., 2020). For variants with heterogeneous effect sizes per ancestry, other local ancestry-aware regression methods could potentially improve the power of detection in admixed populations (Atkinson et al., 2021). Second, the observations in this study that admixed populations harbor a greater power of discovery in GWAS than the ancestral populations is credited to the existence of ancestry variance, independent of specific demographic history of either the admixed or the ancestral population. It is possible that the demographic details or specific assumptions of the genetic architecture would affect the absolute value of power estimate on a finer scale, which has not been the focus of this study, yet is worth being further explored through forward or coalescent simulations with additional details (including modeling differential linkage disequilibrium patterns) in the future. Third, we focused on a single admixture scenario, albeit one reflecting a realistic scenario. We would anticipate our observed patterns to be exaggerated in populations with even contributions from Populations 1 and 2. Further our framework could be extended to k-way admixed populations, but the interpretation and degree of population-specific interpretation become far more complex to be described here.

Despite the underrepresentation of admixed groups in large GWAS, recent research has highlighted the importance of conducting genetic research with more diversity. Our work joins burgeoning efforts to quantify the statistical benefits of complex trait studies in diverse populations, especially populations of mixed ancestry. Our work suggests another advantage for conducting genetic studies in admixed populations, which comes from elevated allele frequencies when traits are moderately to highly differentiated. Moreover, discoveries from such studies aid improvement in cross-population PRS, which is critical in clinical prediction in personalized medicine yet presently has suboptimal performance for many biomedical traits in non-European populations (Martin et al., 2019; Rosenberg et al., 2019; Cavazos and Witte, 2021). We therefore highlight that insights gained from admixed populations provide improved and appealing generalizable properties compared to homogeneous populations. As the field increasingly moves towards personalized medicine applications we must be mindful of opportunities to incentivize novel studies and analyses in diverse and, particularly as we highlight here, populations of mixed ancestry.

## Acknowledgements

This research was supported by NIH grants R35GM133531 (to BMH), R01HG010297, U01HG009080/S1 (to NAZ and CRG) and R01HG011345 (to NAZ and CRG). The content is solely the responsibility of the authors and does not necessarily represent the official views of the National Institutes of Health.

## Conflicts of Interest to declare

None.

## Software Availability

*APRICOT*: https://github.com/menglin44/APRICOT

