## Supplementary Materials for "Admixed Populations Improve Power for Variant Discovery and Portability in Genome-wide Association Studies"

Supplementary Notes

### 1. Simulation-based power estimate between a trait and global ancestry

The first simulator we provide in this study builds hypothetical phenotypes in admixed populations using only global ancestries, without involving genotype. The aim is to assess if the sample size is adequate for observing a dichotomous trait by ancestry correlation.

#### 1.1 Simulating individuals' risks of a disease in an admixed population

Incidence rates, the proportion of a population diagnosed with a disease status in a given period of time, are used in this simulation to represent the probability of an individual in a population of ancestral origin getting the disease. Incidence rate is logit transformed to present the additive form used in binary trait modeling, referred to as proxy phenotype (*Y*).

Admixture percentages for a putative number of admixed individuals can be obtained by sampling with replacement from existing empirical estimates of ancestry proportions. The proxy phenotype for an admixed individual is the ancestry-weighted sum of those from the populations of ancestral origin.

$$Y_{admixed} = \sum_{j=1}^{n} Y_j P_j$$

where $n$ is the total number of ancestry components in the admixed population, and $Y_j$ and $P_j$ are the proxy phenotype and global ancestry percentage of the $j$th ancestral population, respectively. The simulated proxy phenotypes in the admixed population are converted back to risk of developing the disease.

#### 1.2 Simulating case/control status (binary phenotypes) for a proposed sample size of admixed individuals

If a liability threshold is provided, the cutoff is used to divide the distribution of disease risks into cases and controls. If that information is unknown, a case/control status is obtained through a binomial trial with the simulated probabilities. Subsequently, a proposed number of cases and controls for the putative association studies are randomly drawn from the pool.

#### 1.3 Testing correlations between phenotype and ancestry

A logistic regression is performed between an individual's binary phenotype and each ancestry component, with corresponding p values recorded.

#### 1.4 Power estimate

Steps 1.1.1 through 1.1.3 are repeated for a given number of iterations (default 1,000). The power of detecting a significant correlation between a binary phenotype and each ancestry component is defined as the proportion of tested logistic regressions that have p values smaller than a canonical significance threshold (0.05).

### 2. Non-genetic contribution to trait in the genotype-mediated simulator

The non-genetic component is treated as the sum of two parts in admixed populations: (1) random environmental variation modeled as Gaussian noise and (2) environmental confounders correlated with ancestry, such as socioeconomic status and education, modeled as ancestry by environment interaction.

For ancestry-homogenous populations, only the first part applies. The environmental effect for individuals in that population is modeled as a normal distribution with variance adjusted to fit the trait's heritability:

$$\varepsilon_i \sim N\left(0, var(\boldsymbol{G})\frac{1-h^2}{h^2}\right)$$

where $\boldsymbol{G}$ is the vector of PRS computed for N individuals in the population and its variance reflects the phenotypic variance explained by genetics. $h^2$ is the heritability of the trait of interest.

In addition to Gaussian environmental noise, the admixed population can also harbor ancestry-by-environment interaction, which is modeled via two optional modes to reflect the different nature by which environmental factors covary with ancestry.
In the first mode, we sample the ancestral environmental noise from population-specific Gaussian distribution for either ancestry and sum the noise proportional to the global ancestry, so that the non-genetic component for individual *i* is

$$e_i \sim s(\varepsilon_i + \varepsilon_{iA}\theta_{iA} + \varepsilon_{iB}\theta_{iB})$$

where *A* and *B* denote the two populations of ancestral origin and $\theta$ is the global ancestry proportion from either in the *i*th individual. *s* serves as a scaling constant to ensure the variance of *e* is adjusted, so that heritability of the trait in the population remains as $h^2$.

$$s = \sqrt{\frac{var(\boldsymbol{G})}{var(\boldsymbol{G}) + var(\boldsymbol{G_A})[E(\boldsymbol{\theta})^2 + var(\boldsymbol{\theta})] + var(\boldsymbol{G_B})[1 - 2E(\boldsymbol{\theta}) + E(\boldsymbol{\theta})^2 + var(\boldsymbol{\theta})]}}$$

In the second mode, global ancestry has a constant, ancestry-specific effect on the phenotypic variance, thus creating a linear effect on the phenotype.

$$e_i' = s'(\varepsilon_i + \beta_\theta \theta_i)$$

To align the arbitrary $\beta_\theta$ to a more intuitive concept, we introduce another parameter of constant *p* that stands for the percentage of the environment by ancestry interaction over the overall phenotypic variance. By definition,

$$s'^2 {\beta_\theta}^2 var(\boldsymbol{\theta}) = \frac{p}{h^2} var(\boldsymbol{G})$$

Thus, $\beta_\theta$ can be calculated as

$$\beta_\theta = \sqrt{\frac{p(1-h^2)var(\boldsymbol{G})}{h^2(1-h^2-p)var(\boldsymbol{G})}}$$

The scaling factor *s'* is

$$s' = \sqrt{\frac{1-h^2-p}{1-h^2}}$$

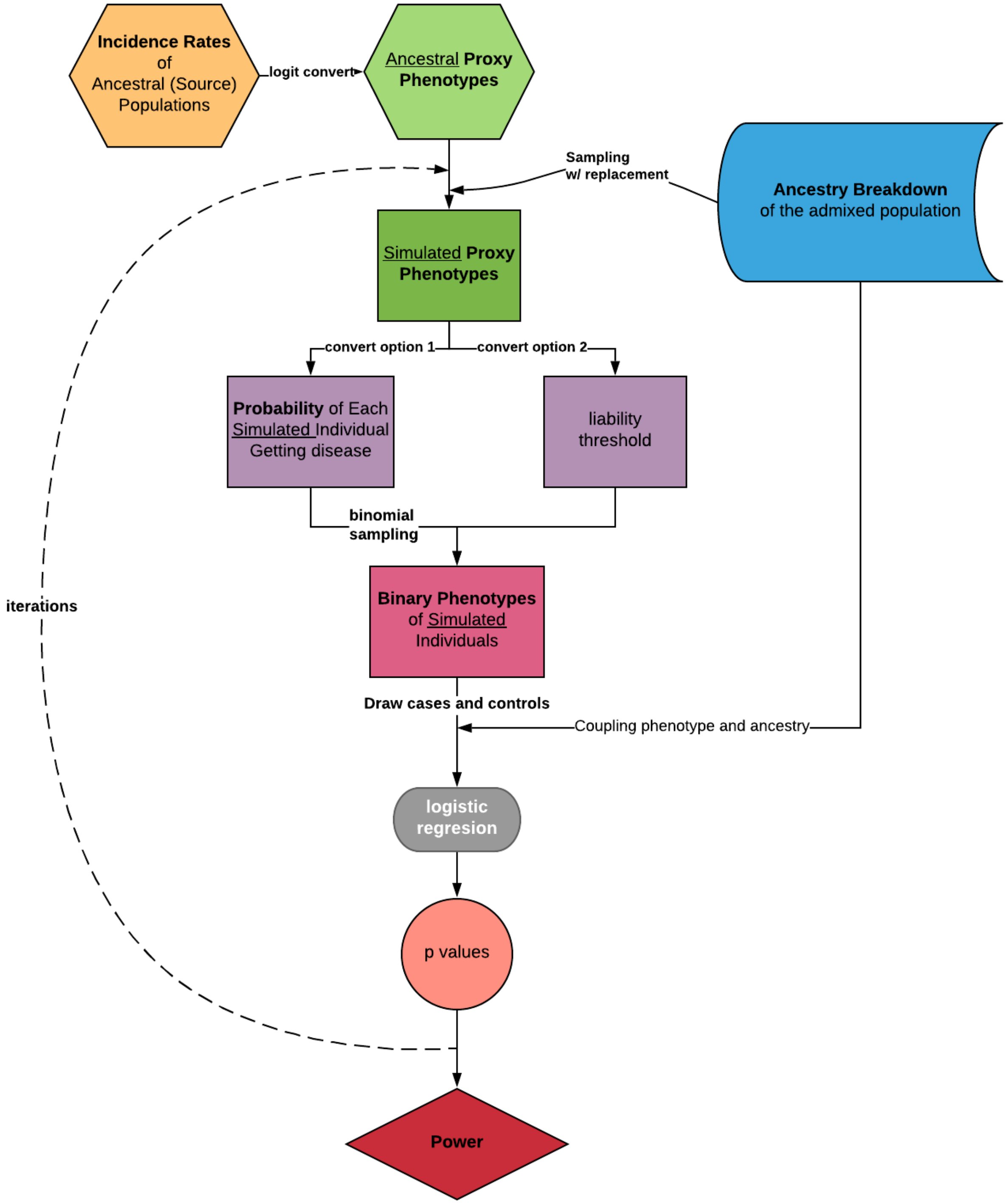


Figure S1. Framework of simulation-based power estimate of correlations between a complex trait and global ancestry.

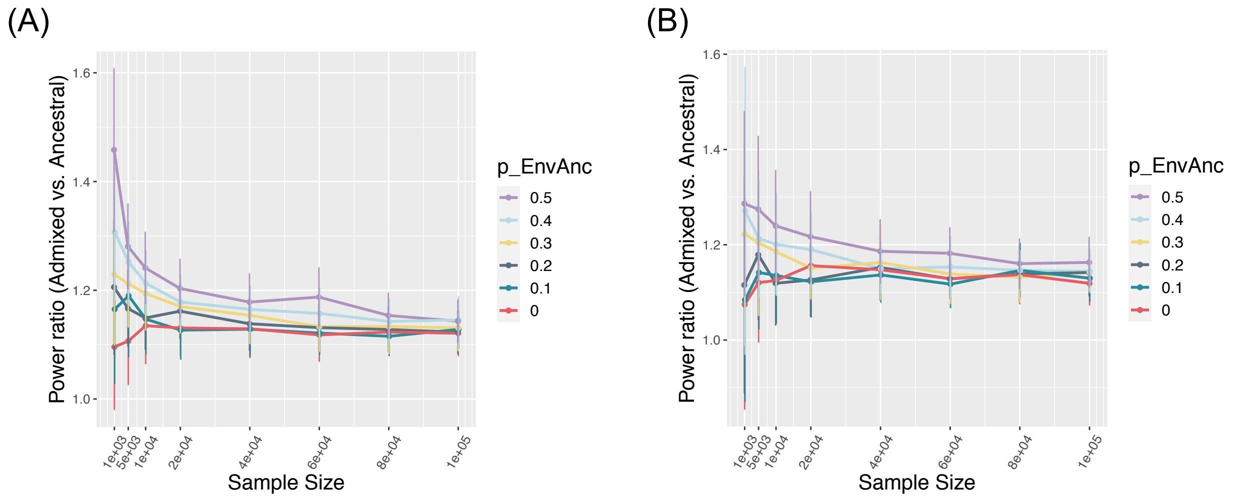


Figure S2. Power ratio when modeling environment by ancestry effect (Env×Anc) as linear dependent on ancestry proportions as in (A) quantitative and (B) dichotomous trait. Other parameters are as the standard setting in the result section. Data point and error bars represent the mean and standard deviation across 50 repetitions, respectively.

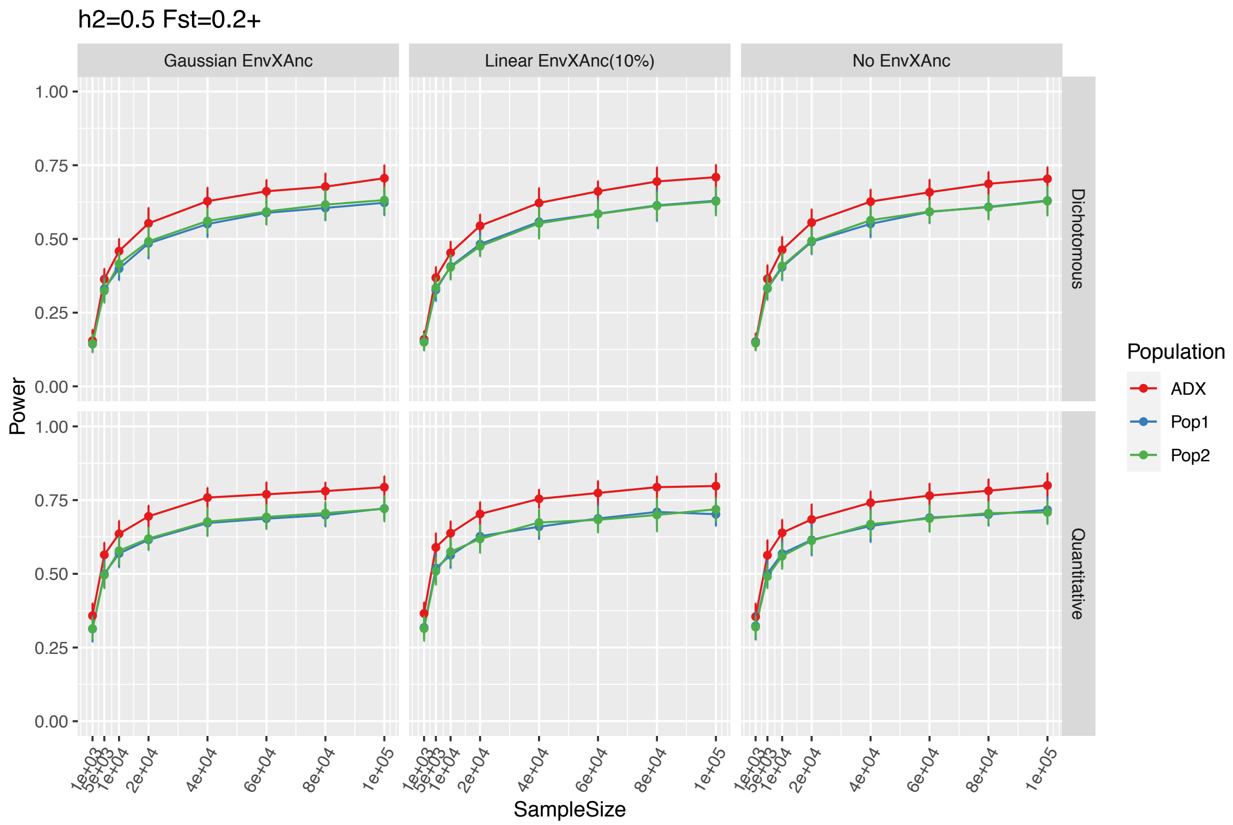


Figure S3. Power of associations in the three putative populations under different modeling of environment by ancestry effect. The simulations are under a setting of heritability 50%, Fst between associated loci as 0.2+, polygenicity of 100 causal variants. Data point and error bars represent the mean and standard deviation across 50 repetitions, respectively.

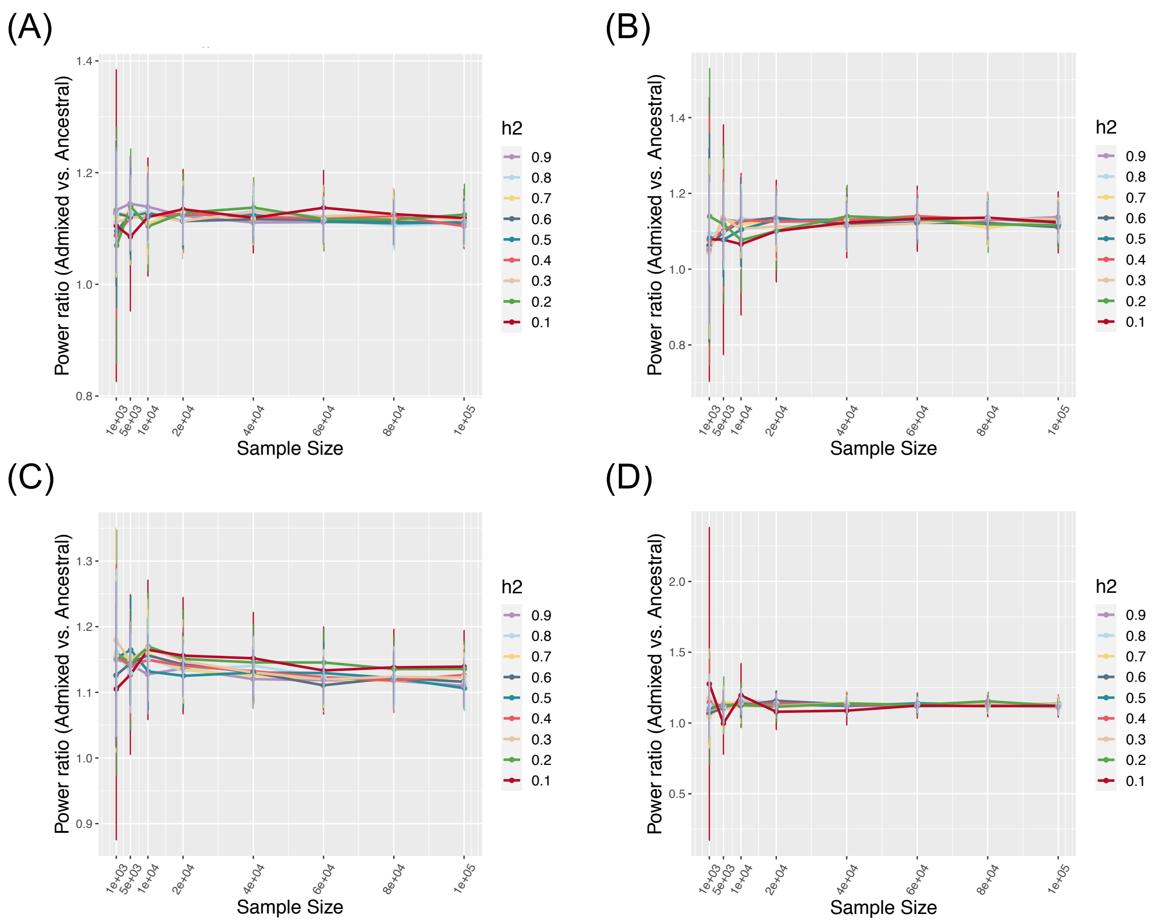


Figure S4. Power ratio over a range of $h^2$ for (A) quantitative trait with Env×Anc modeled as gaussian noise, (B) dichotomous trait with Env×Anc modeled as gaussian noise, (C) quantitative trait with Env×Anc modeled as linear dependent on ancestry, and (D) dichotomous trait with Env×Anc modeled as linear dependent on ancestry. Data point and error bars represent the mean and standard deviation across 50 repetitions, respectively. The other parameters are the same as the standard setting described in the result section.

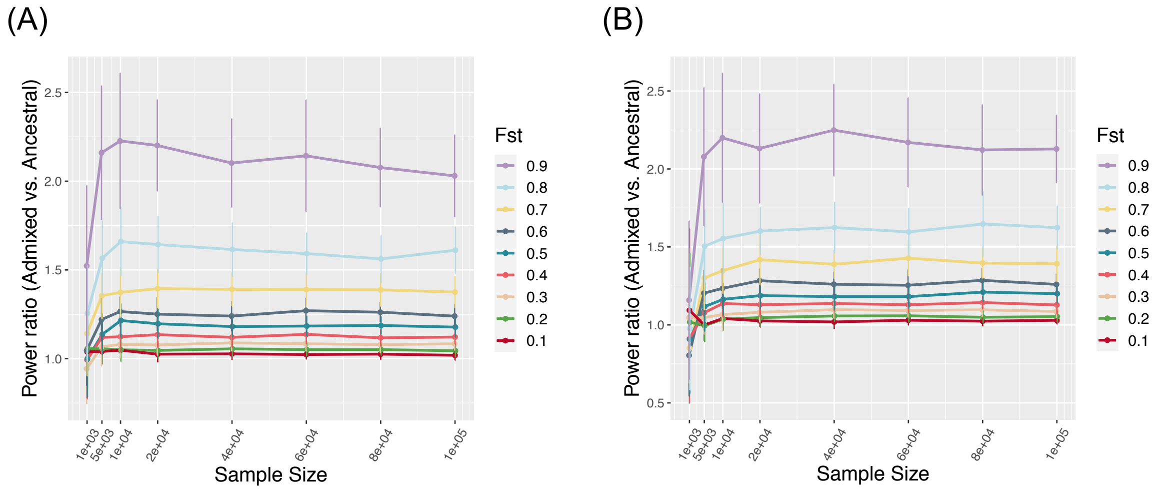


Figure S5. Power ratio over a range of fixed Fst for quantitative trait at a type I error rate of (A) 5e-5 and (B) 5e-8. The other parameters are the same as the standard setting described in the result section. Data point and error bars represent the mean and standard deviation across 50 repetitions, respectively.

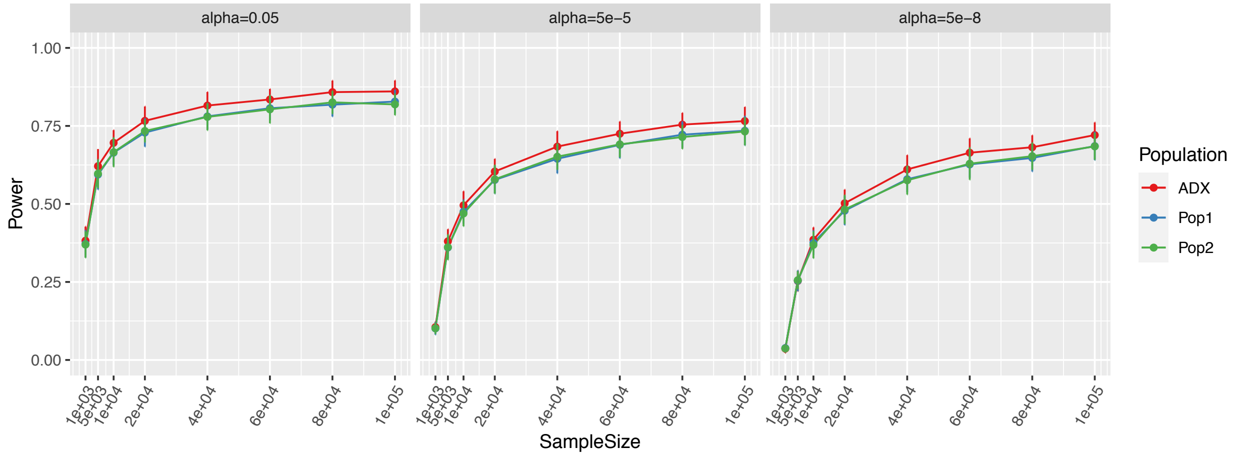


Figure S6. Power of associations at three different type I error rate stringencies. The other parameters are the same as the standard setting described in the result section. Data point and error bars represent the mean and standard deviation across 50 repetitions, respectively.

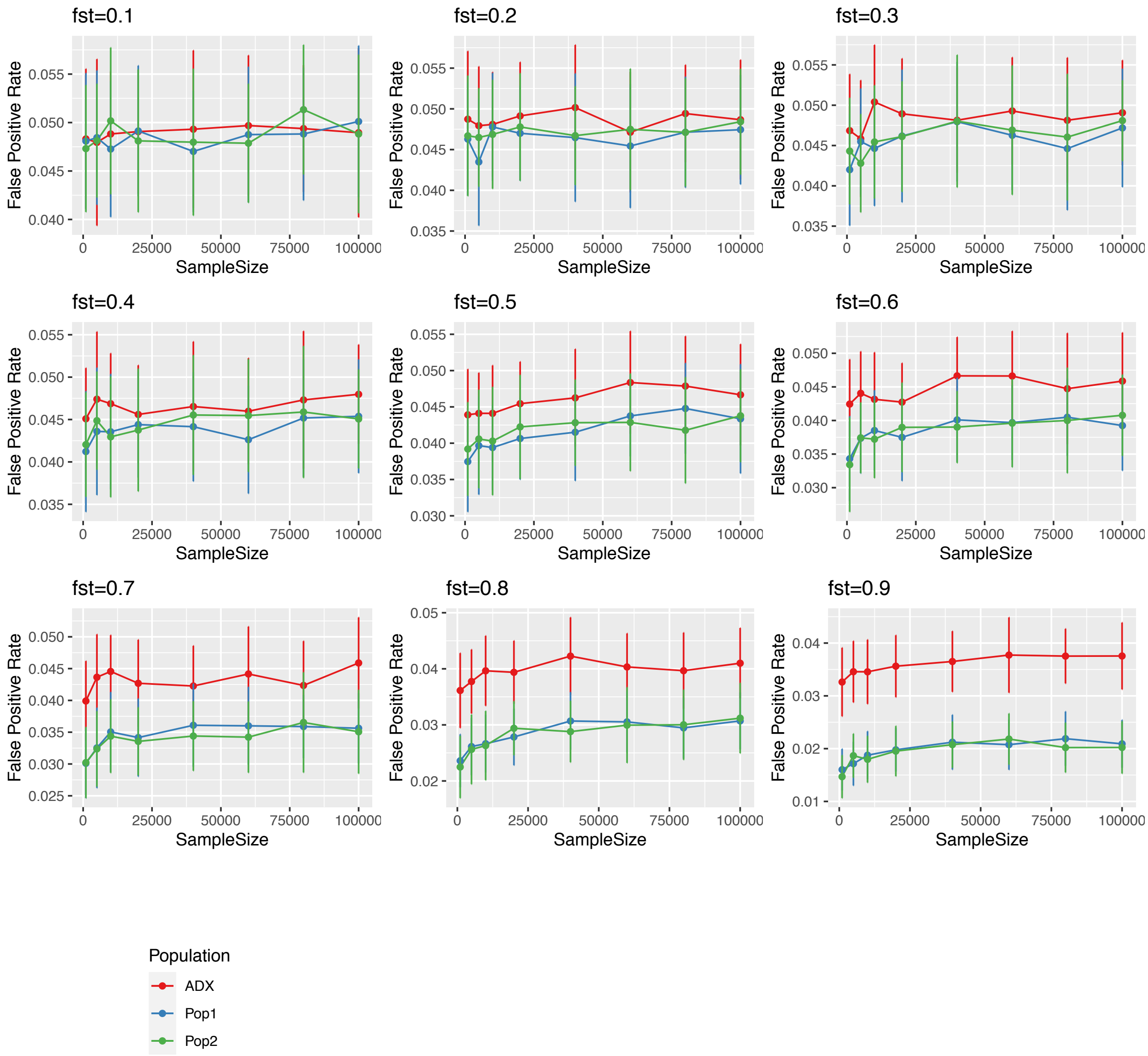


Figure S7. False positive rate assessed in simulations with a range of fixed Fst. The other parameters are the same as the standard setting described in the result section. Data point and error bars represent the mean and standard deviation across 50 repetitions, respectively.

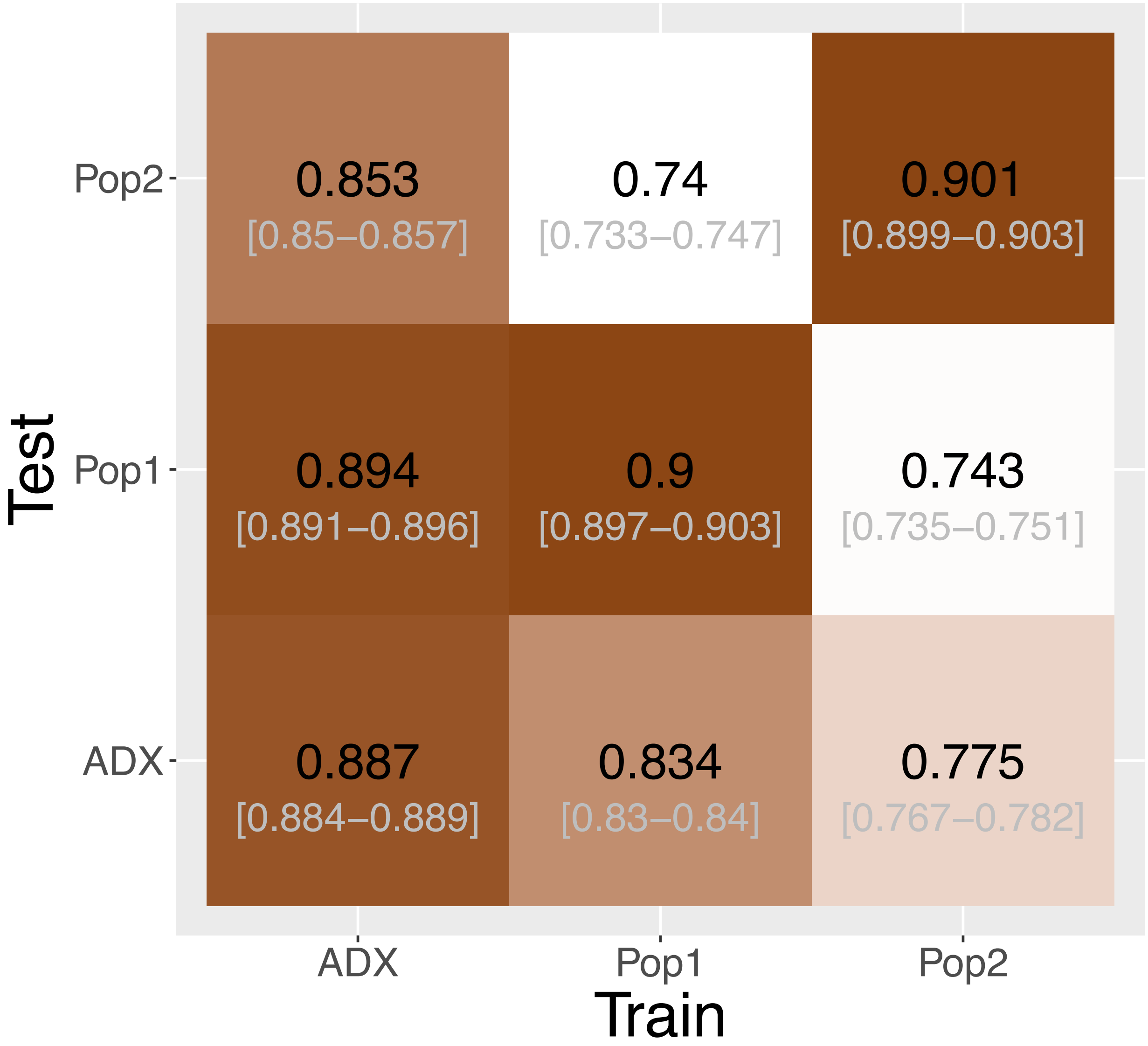


Figure S8. Heatmap of accuracy of PRS using true positive signals at significance level p=0.05. The accuracy is measured as the correlation coefficient between the estimated PRS against the true PRS. The training population where the weights and variants were identified, and the test population in which to construct PRS, are specified on the x- and y-axes. Central numbers in black within each cell are the average correlation coefficient across 50 independent simulations, with the 95% confidence interval of the mean acquired from bootstrapping (n=1,000).

Table S1. Top 50 traits from GWAS catalog with the largest mean $F_{ST}$ at associated variants between CEU and YRI

| Rank | Traits | mean $F_{ST}$[1] |
|---|---|---|
| 1 | Skin pigmentation | 0.620 |
| 2 | Eye color | 0.595 |
| 3 | Skin pigmentation traits | 0.536 |
| 4 | Eye color traits | 0.511 |
| 5 | Hair morphology traits | 0.482 |
| 6 | Skin, hair and eye pigmentation (multivariate analysis) | 0.438 |
| 7 | Heart rate response to beta blockers (atenolol add-on therapy) | 0.404 |
| 8 | Bone ultrasound measurement (broadband ultrasound attenuation) | 0.320 |
| 9 | Urinary metabolites (H-NMR features) | 0.295 |
| 10 | Waist-to-hip ratio adjusted for BMI x sex interaction | 0.281 |
| 11 | Glaucoma (primary angle closure) | 0.280 |
| 12 | Bone properties (heel) | 0.271 |
| 13 | Platelet aggregation | 0.270 |
| 14 | Normal facial asymmetry (angle of surface orientation score) | 0.269 |
| 15 | Red blood cell fatty acid levels | 0.268 |
| 16 | Tanning | 0.265 |
| 17 | Pre-treatment viral load in HIV-1 infection | 0.264 |
| 18 | Tumor biomarkers | 0.252 |
| 19 | Pallidum volume | 0.251 |
| 20 | Caudate nucleus volume | 0.247 |
| 21 | Itch intensity from mosquito bite adjusted by bite size | 0.241 |
| 22 | Knee osteoarthritis | 0.240 |
| 23 | Urate levels in overweight individuals | 0.239 |
| 24 | Rosacea symptom severity | 0.238 |
| 25 | Anthropometric traits | 0.238 |
| 26 | Alcoholic chronic pancreatitis | 0.235 |
| 27 | Parental longevity (combined parental attained age, Martingale residuals) | 0.235 |
| 28 | Coronary artery disease or ischemic stroke | 0.233 |
| 29 | Membranous nephropathy | 0.231 |
| 30 | Mosquito bite size | 0.231 |
| 31 | Putamen volume | 0.228 |
| 32 | Glycerophospholipid levels | 0.227 |
| 33 | Squamous cell carcinoma | 0.225 |
| 34 | Local histogram emphysema pattern | 0.225 |
| 35 | Obesity (early onset extreme) | 0.225 |

| | | |
|---|---|---|
| 36 | Systemic lupus erythematosus and Systemic sclerosis | 0.225 |
| 37 | Longevity | 0.220 |
| 38 | Loneliness | 0.215 |
| 39 | Loneliness (MTAG) | 0.215 |
| 40 | Heart failure | 0.214 |
| 41 | Regular attendance at a pub or social club | 0.214 |
| 42 | Brown vs. black hair color | 0.212 |
| 43 | N-glycan levels | 0.211 |
| 44 | Trans fatty acid levels | 0.209 |
| 45 | Allergic sensitization | 0.208 |
| 46 | Telomere length | 0.203 |
| 47 | Feeling guilty | 0.203 |
| 48 | Cutaneous squamous cell carcinoma | 0.202 |
| 49 | Primary tooth development (number of teeth) | 0.198 |
| 50 | Post bronchodilator FEV1 | 0.198 |

[1]$F_{ST}$ was calculated using Weir and Cockerham's estimate.

Table S2. One-way Wilcoxon P-values of comparing cross-population PRS accuracy

| type I error rate | ADX vs Pop 1 | ADX vs Pop 2 |
|---|---|---|
| Constructed from true positive signals | | |
| 0.05 | 4.54e-18 | 5.13e-18 |
| Constructed from all positive signals | | |
| 0.05 | 2.66e-18 | 6.51e-10 |
| 5e-4 | 3.73e-13 | 6.51e-10 |
| 5e-6 | 5.24e-5 | 4.93e-2 |
| 5e-8 | 0.72 | 0.84 |
